# The Two Tube Volatile Assay: a non-contact benchtop bioassay for monitoring susceptibility to transfluthrin

**DOI:** 10.64898/2026.06.12.731863

**Authors:** Authors: Giorgio Praulins, Amy Lewis, Tim Hill, Boris N’dombidje, Sylvia Milanoi, Gemma Harvey, Daniel P. McDermott, Jeff Jones, Bernard Abong’o, Eric Ochomo, Corine Ngufor, Rosemary Susan Lees

## Abstract

**Introduction:** Progress against malaria has stalled since 2015, with insecticide resistance a key driver. Spatial emanators release volatile insecticides into the air, exposing mosquitoes through a route distinct from the tarsal contact used by treated nets, indoor residual spraying, and standard bioassays. Transfluthrin is currently monitored using the WHO bottle bioassay, which combines contact and vapour exposure and cannot isolate airborne effects. A scalable, vapour-only method is needed to characterise susceptibility to volatile pyrethroids.

**Methods:** We adapted the WHO tube bioassay to deliver non-contact transfluthrin vapour exposure without contacting treated papers. Non-blood-fed female Anopheles were exposed to acetone control or transfluthrin papers at 0.005, 0.1 and 2 mg/paper. Knockdown was recorded over 60 minutes and mortality at 24 hours. Testing spanned three laboratories (LSTM, AIRID and KEMRI) using susceptible reference and resistant lab colonies. Dose response models estimated EC□□ for knockdown (EC□□KD) and mortality (EC□□Mort), with resistance ratios (RR) against the local susceptible reference.

**Results:** The assay generated clear concentration- and time-dependent responses. At LSTM, Kisumu gave an EC□□KD of 0.052 and EC□□Mort of 0.054 mg/paper; Siaya was comparable (RR□□KD 1.17, RR□□Mort 0.89), whereas Tiassalé 13 and KDR showed reduced susceptibility (RR□□KD 2.42 and 2.85; RR□□Mort 5.56 and 6.65). At AIRID, the resistant Cové strain did not reach 50% knockdown or mortality. At KEMRI, Siaya showed reduced susceptibility (RR□□KD 3.36/RR□□Mort 5.65). Kisumu at AIRID and KEMRI was comparable to LSTM (EC□□KD 0.007/EC□□Mort 0.030; EC□□KD 0.044/EC□□Mort 0.023). Despite inter-laboratory variation, susceptible strains remained distinguishable from resistant ones. Lower RRs than for contact pyrethroids suggest contact-based phenotypes may not predict vapour susceptibility.

**Conclusion:** The assay provides a practical benchtop method using standard WHO hardware, distinguishing susceptible from resistant strains across three laboratories. Twenty-four-hour mortality is the recommended endpoint, 60-minute knockdown secondary. Not yet validated, it provides a foundation for further validation and adaptation to other actives.

## Introduction

After major reductions in malaria burden between 2000 and 2015 driven by insecticide-treated nets (ITNs) (2), progress has stalled. The *World Malaria Report 2024* shows that malaria cases have begun to rise again since 2015. The causes are complex, but insecticide resistance is clearly a contributing factor (3).

Spatial repellents have been used in commercial products to reduce nuisance mosquito biting indoors since the 1990s. However, research into their potential as public health tools only expanded in the 2000’s (1), and they received a conditional recommendation from the WHO for deployment for the prevention and control of malaria in August 2025 (4). Spatial emanators offer an alternative route of entry to traditional contact insecticides used in insecticide treated nets (ITNs) and indoor residual spraying (IRS). Rather than requiring tarsal contact, these tools release insecticidal vapour into the air. Mosquitoes entering the treated space may absorb a lethal concentration through respiration or cuticular uptake, or exposure can result in knockdown and other sublethal effects at lower concentrations.

The fact that the effect on mosquitoes is more complicated than the rapid kill induced by exposure to solid-state insecticides has led to discussion over which bioassay endpoints are most appropriate for detecting resistance to volatile pyrethroids. The WHO bottle bioassay (5) is currently recommended for monitoring transfluthrin susceptibility (6). However, because mosquitoes in this assay are exposed both by tarsal contact and to the vapour while confined in a small space, the method may not accurately reflect the route of entry for spatial repellents or capture changes in response which are specifically relevant to resistance developing to volatile pyrethroids. This mismatch may influence both assay outcomes and how results are interpreted in operational contexts. Evidence that these exposure routes can yield distinct outcomes is provided by Kokkas et al. (2026), who demonstrated that transfluthrin response in deli pot assays differed markedly between contact and vapour exposure and that the resistance profile to transfluthrin was not equivalent to that observed for deltamethrin even in strains carrying widely circulating pyrethroid resistance mechanisms (7). This suggests that cross-resistance between contact and volatile pyrethroids cannot be assumed, and that the degree of resistance may differ between the two groups of compounds even though the target-site is shared. This distinction has direct implications for how bioassay results are interpreted in operational contexts: a strain characterised as resistant to a contact pyrethroid may not exhibit the same degree of reduced susceptibility to a volatile formulation, and vice versa, meaning that resistance monitoring programmes relying solely on contact-based assays will lack specificity and may either over- or underestimate the risk to spatial emanator efficacy.

During prospective, cluster-randomised, controlled trial of Mosquito ShieldTM, a transfluthrin-based spatial emanator, in Busia County, western Kenya, testing was carried out in Kenya using the WHO bottle bioassay method for transfluthrin varying from ∼55 to 99% mortality at baseline and ∼75 to 95% mortality during the intervention phase of the trial across 11 sites. Despite the apparent high intensity resistance to transfluthrin in field sampled vectors, spatial emanators based on transfluthrin significantly reduced both the hazard rate of first-time malaria infection and the hazard rate of overall new malaria infections by over 30% (8). This indicates a failure of susceptibility testing using the WHO bottle bioassay to predict the efficacy of products based on transfluthrin.

With WHO having recently endorsed two new spatial repellent emanator products based on transfluthrin (4), and with multiple large-scale trials and pilot deployments of these interventions currently underway (9–11), alongside active development of next-generation volatile actives for emanator use (12), it is timely to critically re-evaluate susceptibility monitoring methodologies for volatile insecticides. An effective assay for this purpose should isolate vapour-phase exposure, avoid tarsal contact, and be capable of capturing toxicity and other relevant effects while remaining simple, reproducible, and scalable across laboratories. It should minimise cross-contamination between test and negative control test units through clear physical separation, appropriate ventilation, and validated cleaning procedures, without reliance on bespoke or highly specialised or costly equipment. In this study, we present results from a bespoke non-contact benchtop bioassay, adapted from the WHO tube bioassay (5). A standard operating procedure (SOP) and protocol for routine susceptibility testing of transfluthrin are provided which can be used to characterise response to transfluthrin in any mosquito population. Adoption of this standardised method will produce data which can be compared between sites and across time in monitoring for changes in susceptibility to transfluthrin because of spatial emanator distribution.

## Materials and Methods

### Mosquitoes

Eight A*n. gambiae* laboratory strains were used to represent susceptible and resistant populations of this species, including parallel colonies of the resistant Siaya strain held independently in two of the sites, and parallel susceptible Kisumu colonies held by all three sites.

For testing at the Liverpool School of Tropical Medicine (LSTM), mosquito colonies were maintained in the LITE facility as described by Williams et al. (15). Insectary conditions were held at 26 ± 2 °C, 70 ± 10% relative humidity, and a 12:12 h light:dark photoperiod with a 1 h simulated dawn and dusk. Larvae were reared in purified water and fed ground TetraMin® tropical flakes (Tetra U.S., Blacksburg, VA, USA). Adults were provided continuous access to a 10% sucrose solution, and females were blood-fed using a Hemotek membrane feeding system (Hemotek Ltd., Blackburn, UK). The blood used for blood feeding was composed of blood plasma and red blood cells supplied by the human blood bank and mixed in a 1:1 ratio upon arrival at LSTM. For the feeding process, a Hemotek membrane feeding system, provided by Hemotek Ltd based in Blackburn, UK, was used.

- The Kisumu strain, originally established from Kisumu, Kenya, has been maintained at LSTM since 1975 and is fully susceptible to insecticides, with no selection pressure applied.
- KDR is a transgenic *An. gambiae* line generated by CRISPR/Cas9 genome editing, referred to as Kisumu-1014F, which carries the kdr target-site mutation (L1014F) in the voltage-gated sodium channel (16). It was produced using the insecticide-susceptible *An. gambiae* Kisumu strain as the genetic background, and the resulting line is homozygous for the introduced mutation. Kisumu-1014F exhibits approximately 14.6-fold resistance to deltamethrin compared with the Kisumu strain.
- The Tiassalé 13 strain, originally established from Côte d’Ivoire, has been maintained at LSTM since 2013 and is routinely selected with a 1-hour exposure to 0.05% deltamethrin (15). Tiassalé 13 shows high resistance to pyrethroids, mediated by both target-site mutations (1014F kdr and ace-1) and enhanced metabolic detoxification via multiple cytochrome P450 enzymes.
- The Siaya strain was originally established from blood-fed indoor-resting females collected in Siaya County, western Kenya, and was first maintained KEMRI from 2020. A parallel colony was established at LSTM in 2023. It shows pyrethroid resistance to deltamethrin and carries both kdr 995F and 995S alleles, with some evidence of cytochrome P450-mediated detoxification based on restoration of susceptibility following PBO pre-exposure.

For testing at AIRID, mosquito colonies were maintained under standard insectary conditions with a 12:12 light: dark photoperiod. Larvae were reared in bowls containing dechlorinated water at a temperature of 27–33°C and relative humidity of 60–95% and were fed daily with a standardized diet (fish food). Adult mosquitoes were maintained in cages at 25–29°C and 60–95% relative humidity, with continuous access to a 10% glucose solution. Females were blood-fed with a live rabbit as required to support colony maintenance and egg production.

- The Kisumu strain – originally established from Kisumu, Kenya, has been maintained locally at AIRID insectary and is fully susceptible to insecticides.
- The Covè strain, a pyrethroid-resistant strain originally established from Covè, southern Benin and maintained at the AIRID laboratories in Cotonou. The strain is composed of a mixture of An. coluzzii and An. gambiae s.s. with the former present at higher frequencies. Recent WHO susceptibility and intensity bioassays have revealed a high intensity of pyrethroid resistance but continued susceptibility to other insecticide classes including chlorfenapyr. Pyrethroid resistance is mediated by a high frequency of the knockdown resistance (kdr) L1014F mutation (87%) and overexpression of metabolic detoxification enzymes particularly CYP6P3.

For testing at KEMRI, mosquito colonies were maintained under standard insectary conditions with a 12:12 h light:dark photoperiod. Larvae were maintained in a larval room at 28 ± 2 °C and reared on Koi Premium fish food (Kaytee Products, Inc., Chilton, WI, USA). Adults were maintained in an adult room at 27 ± 2 °C, with continuous access to a 10% sugar solution provided on cotton wool. Females were blood-fed on goat blood obtained from licenced abattoirs in Kisumu, Kenya using an artificial membrane feeding system (Hemotek Ltd., Blackburn, UK) to support colony maintenance and egg production.

- The Kisumu strain, *Anopheles gambiae* Kisumu, is an insecticide-susceptible reference strain originally established from Kisumu, Kenya. The colony used for testing at KEMRI was established in 2025 from a parallel LSTM colony following contamination of the previous local colony. Kisumu was maintained at 80 ± 10% relative humidity, with no insecticide selection pressure applied.
- The Siaya strain, *Anopheles gambiae* Siaya, is a pyrethroid-resistant strain originally established from mosquitoes collected in Siaya County, western Kenya. The colony was first maintained at KEMRI from 2020, with a parallel colony established at LSTM in 2023. The Siaya colony used for testing at KEMRI was the same colony as that established at LSTM in 2023. Siaya was maintained at 70 ± 10% relative humidity and underwent periodic insecticide selection with deltamethrin every four generations. Resistance is associated with kdr-mediated target-site resistance, with previous characterisation indicating the presence of both kdr 995F and 995S alleles, alongside some evidence of cytochrome P450-mediated detoxification based on restored susceptibility following PBO pre-exposure.

### WHO bottle bioassays

Bottle bioassays using transfluthrin were conducted for all strains included in the study at their respective testing centres to provide comparison with the current standard bioassay recommended for transfluthrin (5). Two control bottles and four treated bottles were tested in accordance with the WHO SOP, using transfluthrin at the discriminating concentration (DC) of 2 µg/bottle with acetone as the solvent. The transfluthrin and acetone used were from the same source as those used in the Two Tube Volatile Assays in each site

### Treatment of filter papers

At LSTM, transfluthrin (Sigma Aldrich, Lot No. BCCN1001; purity: 99.2% and Ehrenstorfer, Lot No. G1379019; purity: 99.11%) was diluted in acetone (JT Baker, Lot No. 2434405854 and Sigma Aldrich, Lot No. SDBB2395) to prepare aliquots of the required test concentrations for application to 12×15 cm filter papers (Whatman, Lot. No18386294) prior to bioassay following the same approach outlined in the (17). Dilutions were adjusted for the purity of the source of transfluthrin given on the Certificate of Analysis.

Based on preliminary data with Kisumu and Tiassalé 13 strains, three concentrations were selected: 0.005, 0.1, and 2 mg/paper. This range of concentrations was chosen to enable the assessment of transfluthrin’s effects on both knockdown and mortality across species and strains. The 0.005 mg concentration represented a low exposure expected to produce no response in Tiassalé 13, the most resistant strain used for method development, and low-level response in Kisumu, the susceptible comparator. The 0.1 mg concentration served as an intermediate level anticipated to capture the differential responses between strains. The 2 mg concentration represented a high exposure expected to cause rapid and substantial knockdown in Kisumu and a readily observable response in Tiassalé 13. KDR was expected to show an intermediate response for each concentration based on previous results (7). By using this range, we expected to be able to characterise the relative response in different mosquito populations, quantified in terms of a resistance ratio relative to a susceptible reference strain.

Equivalent source information was recorded for the additional test sites. At AIRID, transfluthrin was obtained from Envu (Batch No. PMLO000650; purity: and acetone was obtained from Sigma Aldrich). At KEMRI, transfluthrin was obtained from Sigma-Aldrich, USA (Batch No. BCCN1001; purity 99.2%) and acetone was obtained from Griffchem, India (Batch No. AL032517).

### Adaptation of the WHO tube system

The Two Tube volatile assay adapts the standard WHO tube bioassay kit (supplied by Universiti Sains Malaysia, Penang, Malaysia (17)). Two clear tubes are joined by a central gate, with a mesh barrier between them to prevent mosquitoes from making physical contact with the treated substrate. Treated filter paper is placed in one tube; mosquitoes are loaded into the other (Figure 1).

**Figure 1.**
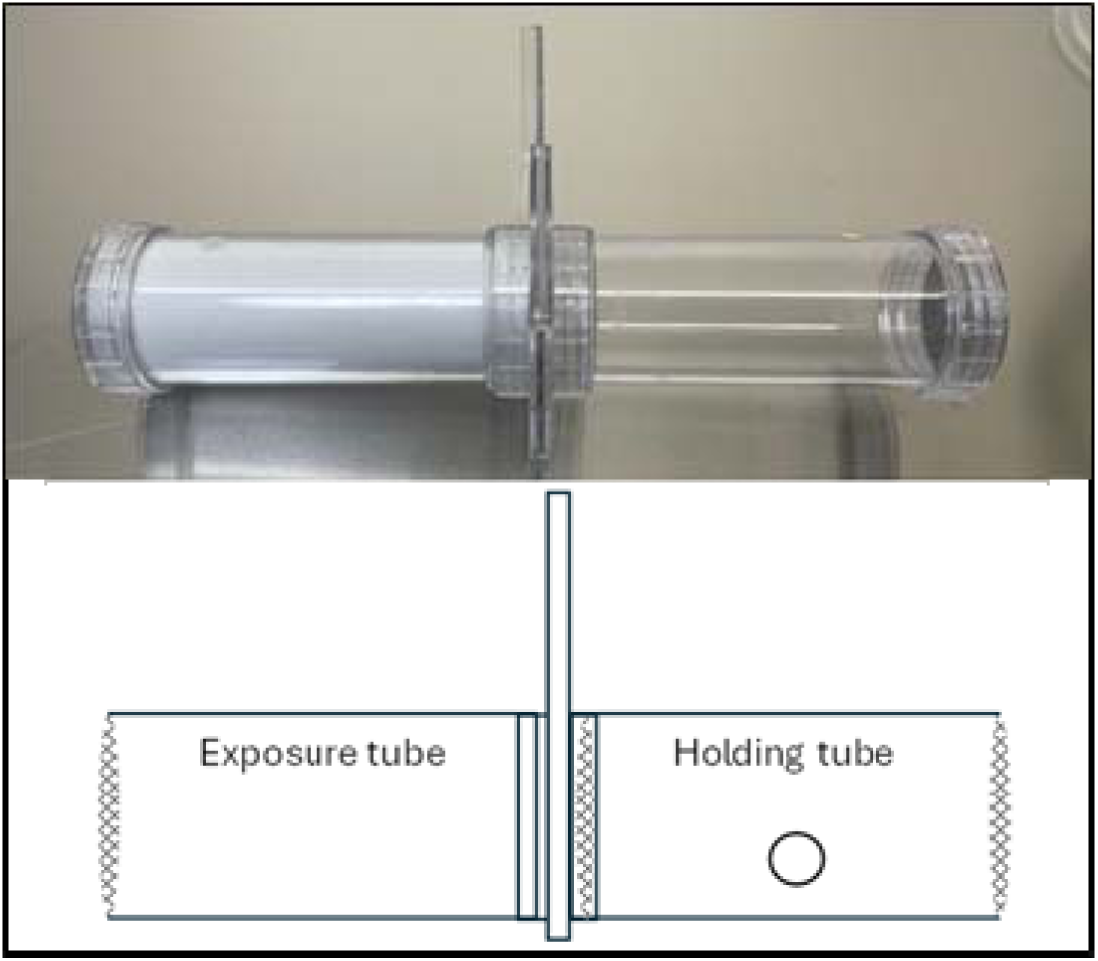
Photograph and schematic diagram of the Two Tube Volatile assay setup, used to evaluate mosquito responses to volatile stimuli without direct contact exposure. (Photo credit: Giorgio Praulins)

When the gate is opened, mosquitoes can move freely between compartments. Cotton wool supports can be used to keep the apparatus level, and the assay is conducted within a fume hood or a well-ventilated area to prevent build-up of volatile compounds.

### Two Tube Volatile Assay method

Mosquitoes were tested in batches of 25 per tube. Each tube was run individually within a fume hood (Faster Chemfast 15, Serial No. 265 and Faster Chemfast 12, Serial No. 453) operating at ∼0.57 m/s airflow, beginning with the negative control (acetone alone), and progressing through increasing concentrations to minimise the risk of cross-contamination

Mosquitoes were aspirated into the holding tube and a small piece of parafilm was used to cover the hole making sure not to obscure the view within the tube. Mosquitoes were allowed to acclimatise for 1 hour before exposure. A 12 × 15 cm sheet of Grade 1 Whatman filter paper (Whatman, Lot No. 18386394) was placed on a wire rack to support it during treatment while minimising transfer of insecticide to the surface below. A 2 mL aliquot of either negative control or insecticide solution was applied evenly to the paper in two 1 mL portions, allowing partial drying between applications to prevent oversaturation. The dilution bottle was then rinsed with a further 1 mL (12).

Once visibly dry, (approximately 1 minute) the treated paper was inserted into the exposure tube, which was then attached to the holding tube and placed horizontally in the fume hood. Cotton wool supports were placed under each end of the tube assembly to keep it level. Opening the slider door marked the start of the assay.

Knockdown was scored every 5 minutes for the first 15 minutes and then every 15 minutes for the remaining 45 minutes for a 1-hour total exposure time. The negative control was only scored every 15 minutes as no knockdown activity was expected in the first 15 minutes.

Knockdown was scored according to the definition described in the WHO “Standard operating procedure for testing insecticide susceptibility of adult mosquitoes in WHO tube tests” as any mosquito unable to stand or fly in a coordinated manner at the scoring timepoint. These included mosquitoes showing no sign of life, those unable to stand properly, those lying on their back and moving their legs or wings but unable to take off, and those able to stand or take off only briefly before falling. Mosquitoes were considered alive only if they could both stand and fly in a coordinated manner (17).

A WHO holding tube and its paired exposure tube are collectively referred to as a test unit. Test units using mosquitoes from the same cohort were considered technical replicates, while those using mosquitoes from different cohorts were considered biological replicates.

For each site, dose, and strain combination, assays were performed using replicate tubes containing approximately 25 adult female mosquitoes per tube. At LSTM, three independent replicates were conducted across three separate assay days, with one replicate performed per day. At KEMRI and AIRID, three replicate tubes were tested for each strain and dose on the same assay day. The AIRID and KEMRI datasets therefore provide within-day replicate data, while the LSTM dataset captures both within-day assay variation and between-day variation.

The sample size was defined as the total number of mosquitoes contributing to the dataset for each site and strain. AIRID and KEMRI each contributed 300 mosquitoes per strain, comprising three replicate tubes per dose across four doses. At LSTM, total sample sizes ranged from 221 to 229 mosquitoes per strain because of minor variation in the number of mosquitoes tested per replicate tube. The LSTM sample sizes were 221 mosquitoes for Kisumu, 223 for KDR, 228 for Siaya, and 229 for Tiassalé 13.

### Cleaning

Across all three sites, bioassay equipment (WHO tubes and associated apparatus) was decontaminated after each use to minimise residual insecticide and cross-contamination between experiments, with the specific reagents and durations varying by site according to chemical availability.

At LSTM, equipment was disassembled and soaked overnight in 5% Decon90 (Decon Labs Ltd), thoroughly rinsed with deionised water, and dried in a drying cabinet (LEEC Drying Cabinet F2, Serial No. 5076) until completely dry. Between biological replicates, fume hoods were wiped down internally with 5% Decon90 to prevent transfluthrin buildup, rinsed twice with deionised water (Elix Essential 10UV water purification system, Serial No. F7CA82554 B), and given a final wipe with 70% ethanol (Severn Biotech, Lot No. 24486).

At AIRID, WHO tubes were cleaned following standard laboratory procedures to prevent cross-contamination. Tubes were first rinsed thoroughly with clean water to remove any visible residues. They were then washed with a mild laboratory detergent, rinsed again with clean water, and allowed to air dry completely before reuse. When required, additional rinsing with acetone was performed to ensure removal of any residual insecticide. All equipment was fully dried prior to subsequent use.

At KEMRI, one of two methods was used depending on chemical availability. In the primary method, tubes were immersed in 5% Decon90 (Decon Labs Ltd) for 3 hours, rinsed thoroughly under running distilled water, and air-dried at room temperature in a dust-free environment. When Decon90 was unavailable, tubes were soaked overnight in distilled water containing liquid soap detergent (Pride® Multipurpose Liquid detergent), rinsed multiple times with distilled water to remove soap residue, given a final rinse with 70% ethanol (Scharlau, Scharlab S.L, Lot No. ET00052500), and air-dried at room temperature. Tubes were visually inspected for cleanliness, cracks, or chips before storage in clean, dust-free containers.

In all cases, equipment was fully dried before subsequent use.

### Statistical analysis

All analyses were conducted in R, using the packages readxl, dplyr, stringr, tidyr, drc and ggplot2.

Knockdown was expressed as the proportion of mosquitoes knocked down at each timepoint and summarised by colony and concentration. For each colony and concentration, individual replicate knockdown trajectories were plotted over time, together with the mean knockdown across replicates.

Twenty-four-hour mortality was calculated as the proportion of mosquitoes dead at 24 hours and summarised by colony and concentration and was likewise plotted as individual replicate values with the across-replicate mean.

To estimate EC_50_ values, dose-response models were fitted for 60-minute knockdown and 24-hour mortality. Each endpoint was analysed as a binomial outcome, with the number affected relative to the number exposed, using a two-parameter log-logistic model (LL.2), which assumes lower and upper asymptotes fixed at 0 and 1, respectively. Dose was treated as a continuous variable, and models were estimated by maximum likelihood. Replicate-level observations were weighted by the number of mosquitoes tested. The zero-concentration control was excluded from EC_50_ fitting, as the log-logistic model is defined on the log-dose scale and cannot accommodate a dose of zero.

For each laboratory site and endpoint, strain-specific curves were fitted within a joint LL.2 model using strain as the curve identifier. Analyses were restricted to strains with at least three unique non-zero dose levels. EC_50_ values and their 95% confidence intervals were derived from the fitted models using the delta method. These are referred to throughout as EC_50_KD, the effective concentration at which 50% of exposed mosquitoes are knocked down at 60 minutes, and EC_50_Mort, the effective concentration at which 50% of exposed mosquitoes die within 24 hours.

All valid observations were retained for model fitting. However, EC_50_ values were only reported where the response reached the 50% effect threshold within the tested dose range, specifically, where every replicate reached at least 50% response at the highest tested concentration and the modelled EC_50_ fell at or below that concentration. Where these conditions were not met, model-derived estimates were retained for audit but excluded from reporting because they required extrapolation beyond the supported response range.

Resistance ratios were estimated within the same joint model for each laboratory site and endpoint, as the model-based ratio of each strain’s EC_50_ to that of the susceptible Kisumu reference strain, with 95% confidence intervals obtained by the delta method (drc::EDcomp). These are referred to as RR_50_KD and RR_50_Mort for knockdown and mortality, respectively, and were reported only where both the test and reference strains met the reporting threshold above.

## Results

### WHO bottle bioassay

All three colonies of Kisumu were defined as susceptible to transfluthrin (>98% mortality 24 hours after exposure) through application of the discriminating concentration (DC) or transfluthrin in a WHO bottle bioassay. All other colonies included in this study were defined as resistant to transfluthrin in this same assay (≤98% mortality) (Figure 2).

**Figure 2.**
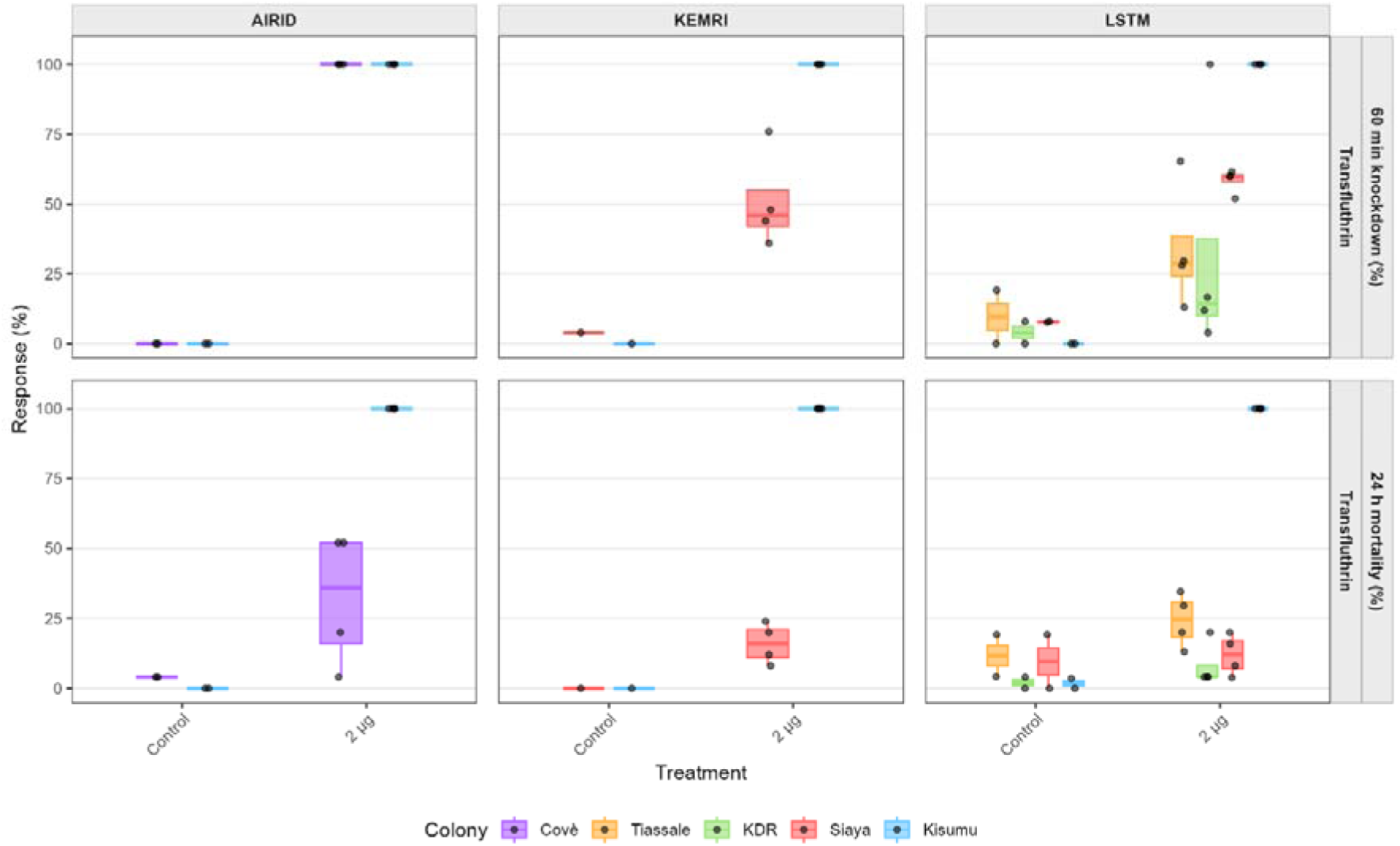
Knockdown and twenty-four-hour mortality of Anopheles gambiae strains exposed to transfluthrin in the WHO bottle bioassay. Mosquitoes were exposed to transfluthrin at 0 and 2 µg/bottle. Panels are facetted by site and endpoint, showing 60-minute knockdown and 24-hour mortality for strains tested at KEMRI, AIRID and LSTM. Boxplots show replicate variation for each colony at each concentration. Boxes represent the interquartile range, horizontal lines show the median, whiskers show the spread of values within 1.5× the interquartile range, and overlaid points show individual replicate values.

At AIRID, the Cové strain showed moderate 24-hour mortality (∼35–50%) at 2 µg/bottle, but negligible knockdown. At KEMRI, the Siaya strain showed moderate 60-minute knockdown (∼50%) at the same concentration, although 24-hour mortality was lower (∼15%). At LSTM, the Tiassalé, KDR, and Siaya strains all exhibited modest knockdown (∼25–35%) at 2 µg/bottle; however, 24-hour mortality remained low across all strains (∼15%).

### Two Tube Volatile Assay

Testing was first performed with four strains of An. gambiae at the Liverpool School of Tropical Medicine (LSTM). For each strain and dose combination, three independent replicates were conducted across three separate assay days, with one replicate per day and approximately 25 adult female mosquitoes per tube. Total sample sizes were 221 mosquitoes for Kisumu, 223 for KDR, 228 for Siaya, and 229 for Tiassalé 13. Knockdown increased with both concentration and time in all strains tested, but the rate and extent of response differed between strains (Figure 3). Kisumu showed the fastest and greatest knockdown response overall, while Tiassalé 13 and KDR were less affected and showed greater evidence of reduced susceptibility. Siaya was closer to the susceptible Kisumu reference strain than to the more resistant strains. The clearest separation between strains was observed at 0.1 mg/paper, while 2 mg/paper produced high or complete knockdown in all strains within the 60-minute exposure period.

**Figure 3.**
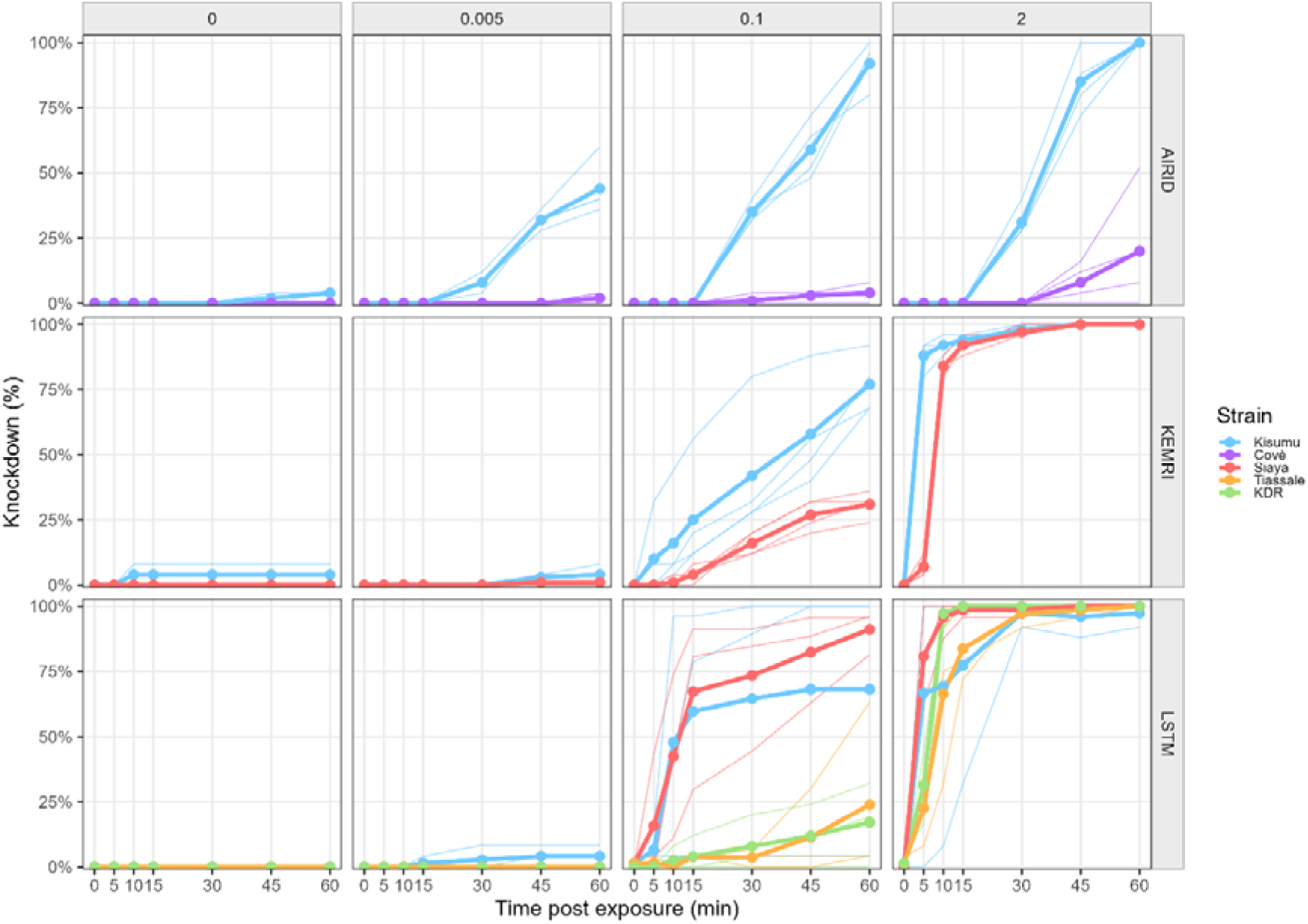
Knockdown in Two Tube volatile assays across LSTM, AIRID and KEMRI. Adult female mosquitoes were exposed to transfluthrin-treated papers at 0, 0.005, 0.1 or 2 mg per paper, with knockdown recorded at 0, 5, 10, 15, 30, 45 and 60 minutes post exposure. Assays at LSTM used Kisumu, KDR and Tiassalé 13; assays at AIRID used Kisumu and Cové; and assays at KEMRI used Kisumu and Siaya. Plot columns represent transfluthrin dose and rows represent testing site. Each strain and dose combination was tested using three replicate tubes containing approximately 25 females each. Points and overlaid lines show mean knockdown percentage over time, while pale replicate lines indicate variation between replicate tubes.

A similar pattern was observed for 24-hour mortality (Figure 4). Kisumu and Siaya showed high mortality at the upper end of the dose range, while Tiassalé 13 and KDR were less affected across the lower concentrations and required the highest concentration to produce strong mortality responses.

**Figure 4.**
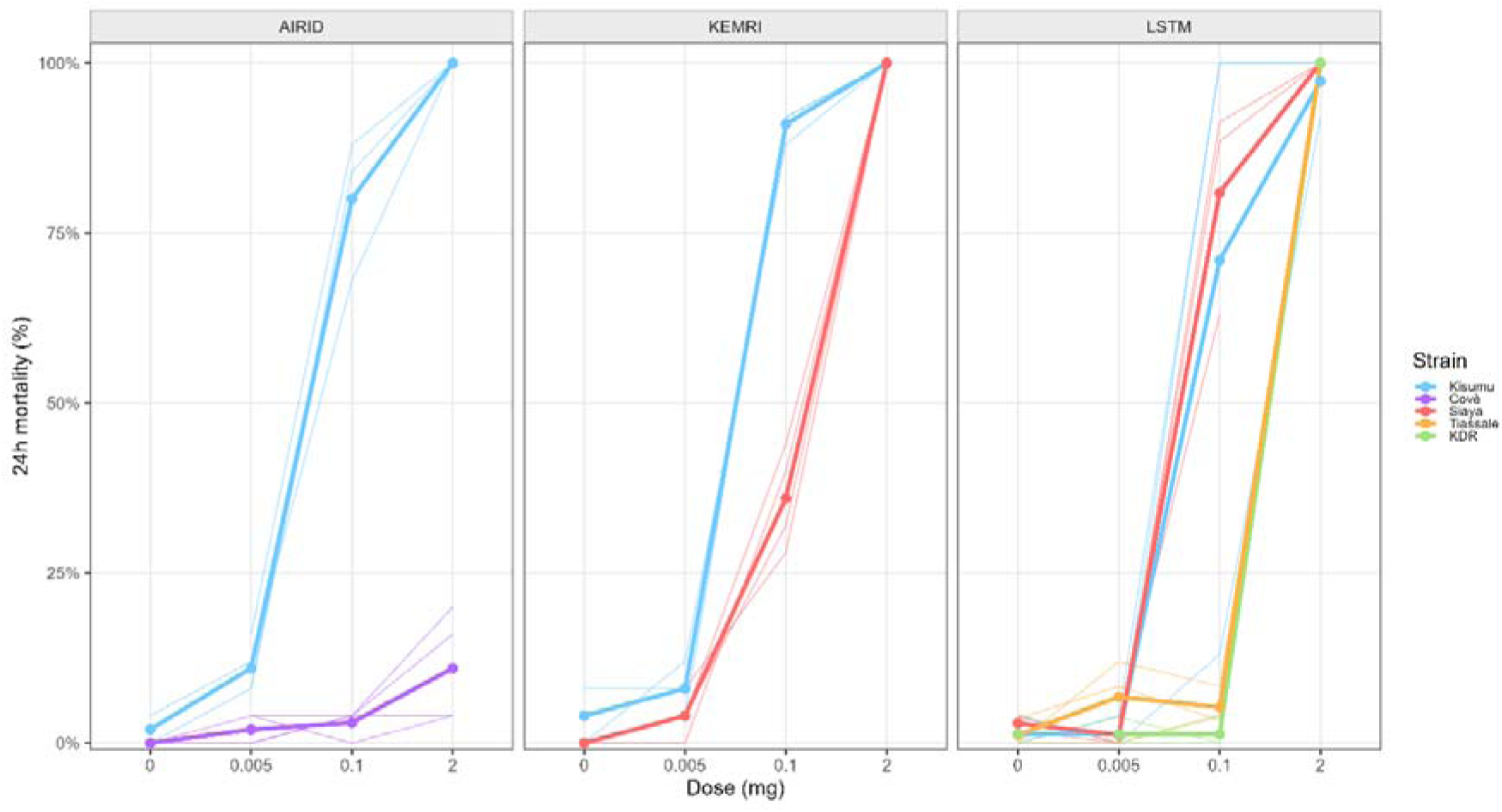
Twenty-four-hour mortality following Two Tube volatile assays across LSTM, AIRID and KEMRI. Adult female Anopheles gambiae sensu lato were exposed to transfluthrin-treated papers at 0, 0.005, 0.1 or 2 mg per paper, with mortality recorded 24 hours after exposure. Assays at LSTM used Kisumu, KDR and Tiassalé 13; assays at AIRID used Kisumu and Cové; and assays at KEMRI used Kisumu and Siaya. Plot panels represent testing site. Each strain and dose combination was tested using three replicate tubes containing approximately 25 females each. Points and overlaid lines show mean 24-hour mortality percentage across replicates, while pale replicate lines indicate variation between replicate tubes.

**Figure 5.**
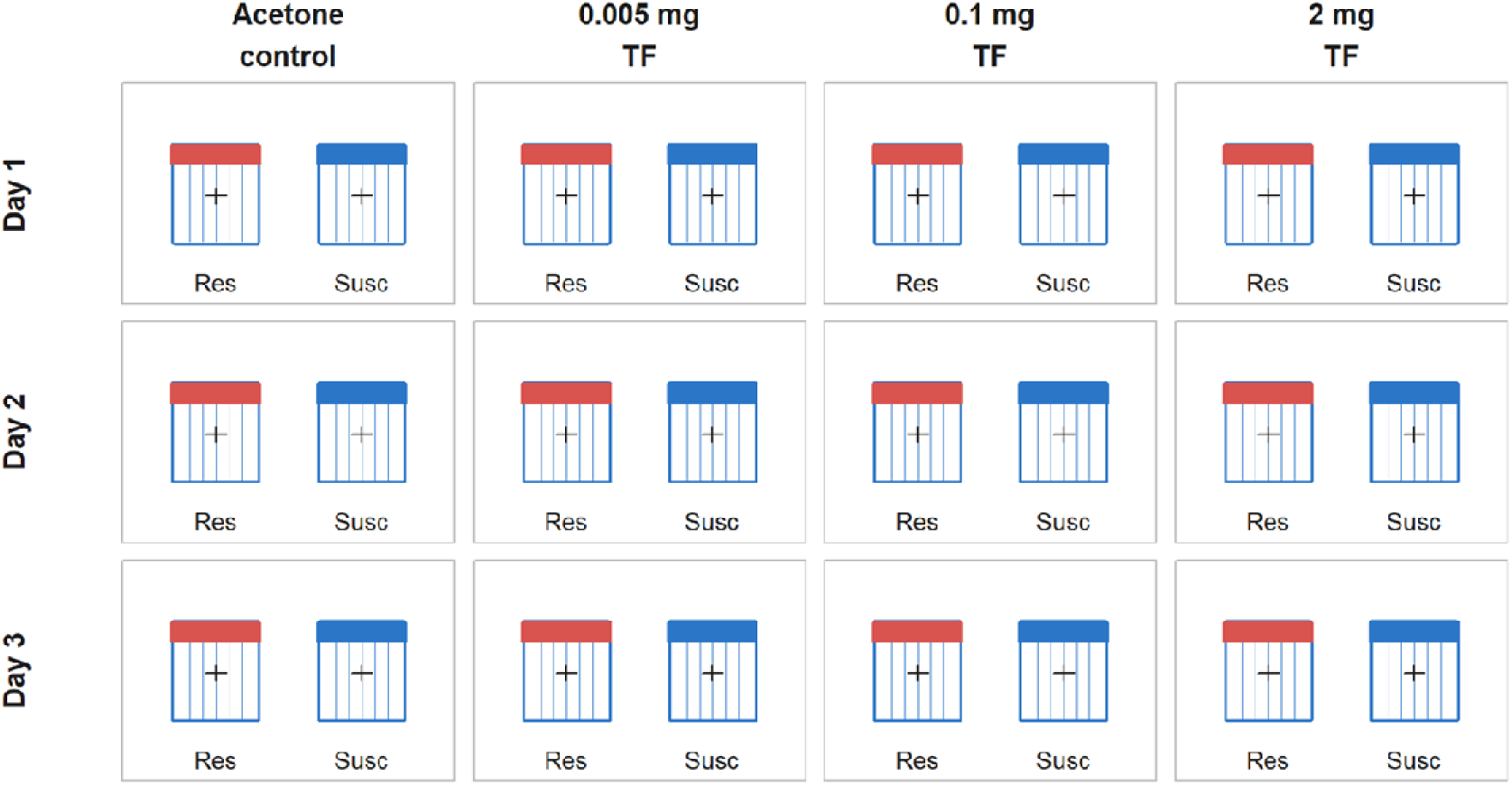
Schematic overview of the routine screening design for the Two Tube Volatile Assay. On each experimental day, a susceptible reference strain (Susc, blue) and a field/resistant strain (Res, red) are tested in parallel under identical conditions. Four treatments are run concurrently: acetone control, 0.005 mg transfluthrin, 0.1 mg transfluthrin and 2 mg transfluthrin per paper. The layout is repeated across three independent experimental days to provide biological replication. This design enables direct comparison of knockdown and 24-hour mortality responses between resistant and susceptible strains at each concentration under standardised vapour only exposure.

Results generated at AIRID also showed time and concentration-dependent responses, with clear separation between the resistant Cové strain and the susceptible Kisumu reference strain. At this site, three replicate tubes per strain and dose were tested on the same assay day, each containing approximately 25 adult female mosquitoes, giving 300 mosquitoes per strain in total. Knockdown was slower than at LSTM, and Cové did not reach 50% knockdown or 50% mortality within the tested dose range, including at the highest dose of 2 mg/paper. In KEMRI, dose-dependent knockdown and mortality responses were similarly observed across strains, with three replicate tubes tested per strain and dose on the same assay day, giving 300 mosquitoes per strain in total. Clear separation between the Siaya field strain and the Kisumu reference was apparent across both knockdown and mortality endpoints (Figure 3).

Mosquito control programmes monitoring susceptibility to insecticides require simple metrics to classify susceptibility and support decisions about the deployment of vector control tools. Dose-response data were therefore fitted to log-logistic dose-response models to estimate the concentration at which 50% of exposed mosquitoes were knocked down after 60 minutes, EC□□KD, or killed after 24 hours, EC□□Mort. Resistance ratios, RR□□KD and RR□□Mort, were then calculated relative to the susceptible reference strain at each site to describe shifts in response between mosquito populations and to support future monitoring over time (Table 1 - Dose response estimates for transfluthrin induced knockdown and twenty-four-hour mortality in Anopheles gambiae s.l. colonies tested across AIRID, KEMRI and LSTM. Estimated doses are shown for the concentration of transfluthrin, expressed as mg per treated paper, required to produce a 50% response for each endpoint. Knockdown estimates refer to 60-minute knockdown, while mortality estimates refer to mortality recorded 24 hours after exposure. Estimates are presented with standard errors and lower and upper 95% confidence intervals. Resistance ratios were calculated relative to the susceptible Kisumu comparator within each site where appropriate. Colonies were excluded from analysis where the fitted response did not reach the 50% effect threshold, preventing reliable estimation of the endpoint.Table 1)Table 2.

**Table 1.**
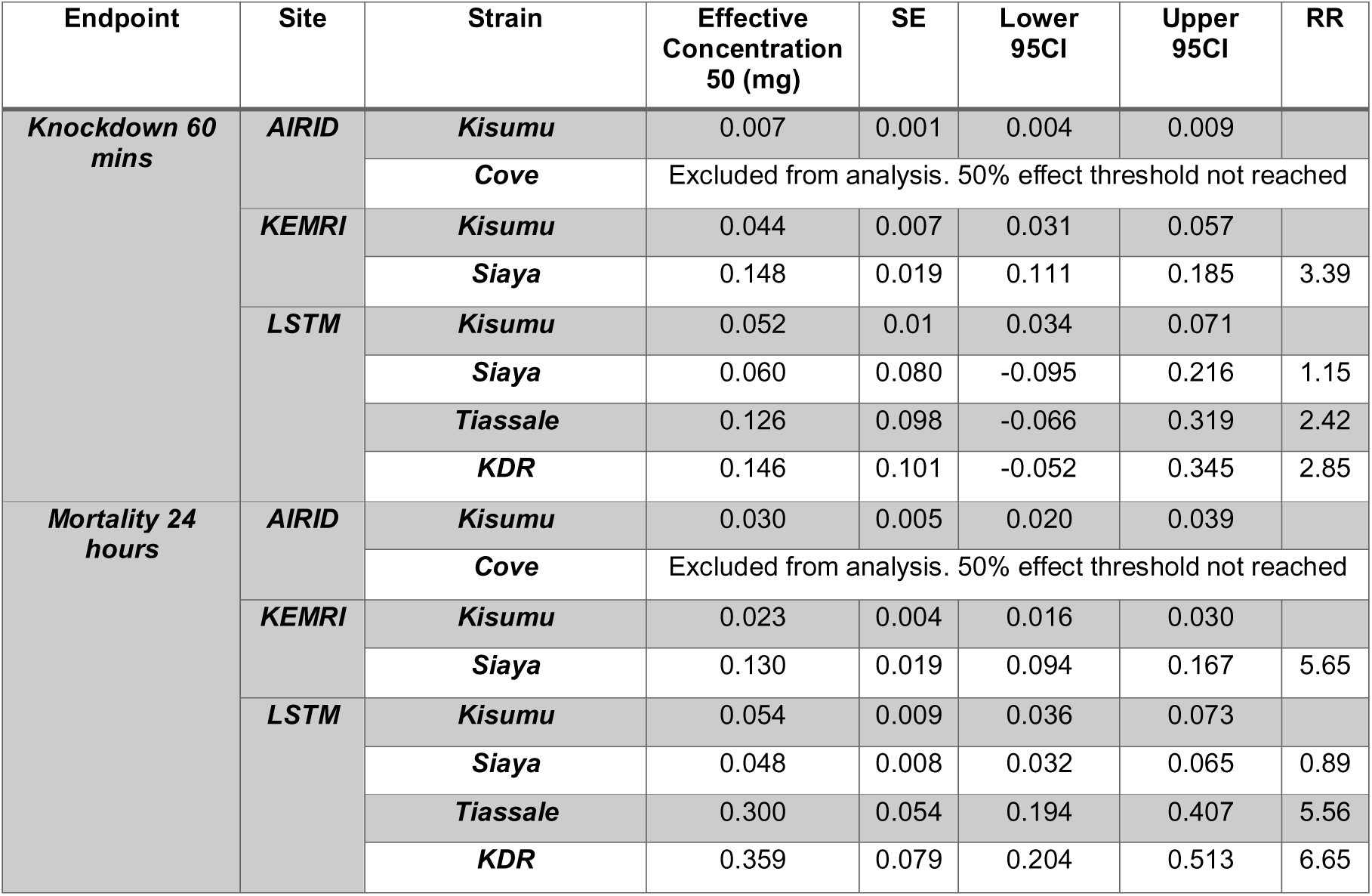
Dose response estimates for transfluthrin induced knockdown and twenty-four-hour mortality in Anopheles gambiae s.l. colonies tested across AIRID, KEMRI and LSTM. Estimated doses are shown for the concentration of transfluthrin, expressed as mg per treated paper, required to produce a 50% response for each endpoint. Knockdown estimates refer to 60-minute knockdown, while mortality estimates refer to mortality recorded 24 hours after exposure. Estimates are presented with standard errors and lower and upper 95% confidence intervals. Resistance ratios were calculated relative to the susceptible Kisumu comparator within each site where appropriate. Colonies were excluded from analysis where the fitted response did not reach the 50% effect threshold, preventing reliable estimation of the endpoint.

**Table 2.**
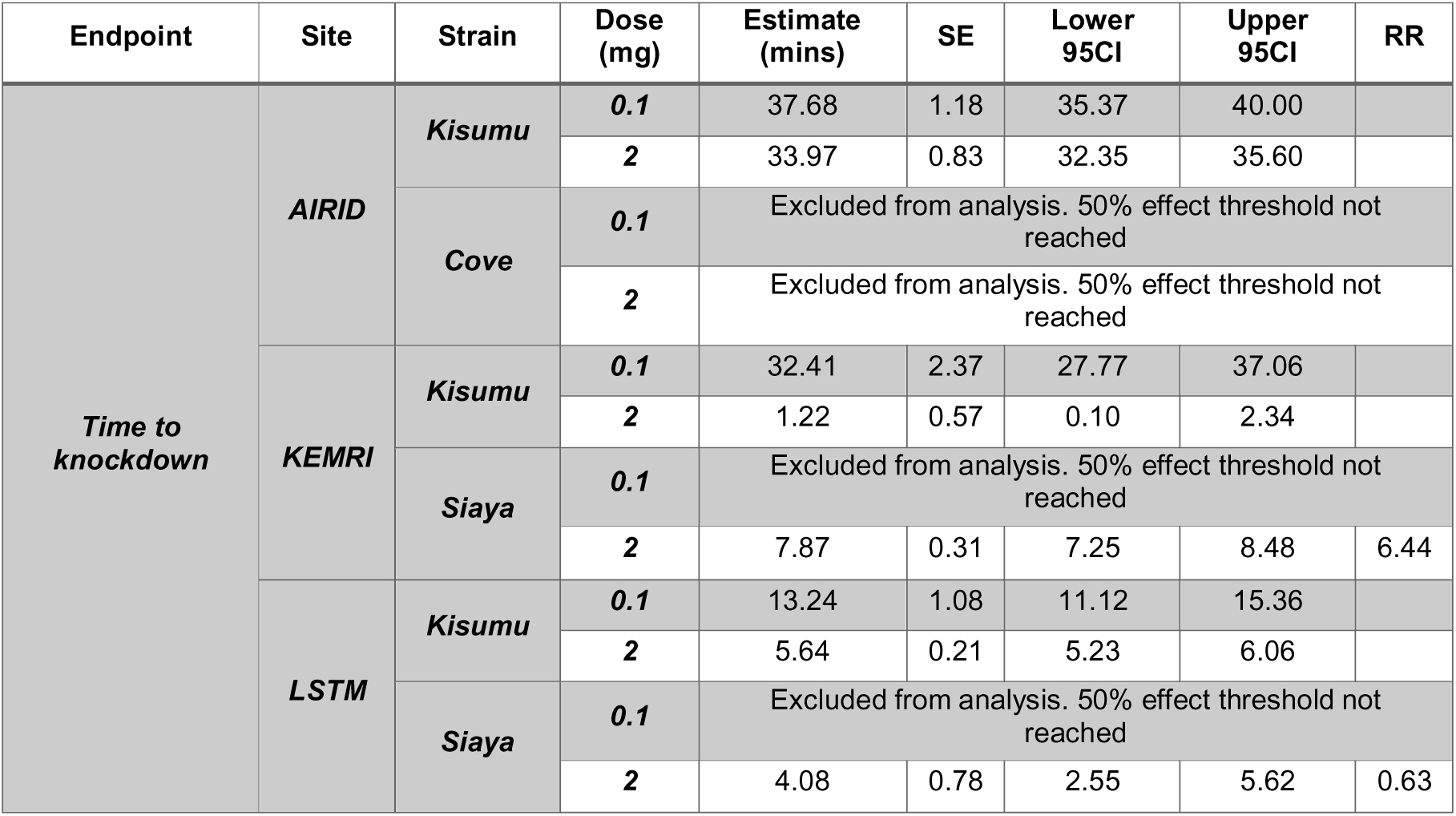
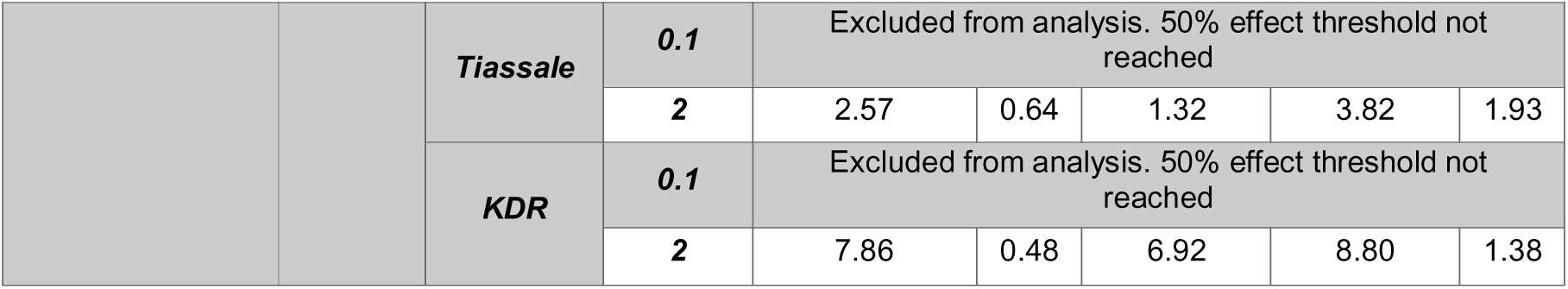
Time to knockdown estimates for transfluthrin exposed Anopheles gambiae s.l. colonies tested across AIRID, KEMRI and LSTM. Estimated time to 50% knockdown is shown for each colony following exposure to transfluthrin treated papers at 0.1 and 2 mg per paper. Estimates are expressed in minutes and are presented with standard errors and lower and upper 95% confidence intervals. Resistance ratios were calculated relative to the susceptible Kisumu comparator within each site and dose where appropriate. Colonies were excluded from analysis where the fitted knockdown response did not reach the 50% effect threshold, preventing reliable estimation of time to 50% knockdown.

| Endpoint | Site | Strain | Dose (mg) | Estimate (mins) | SE | Lower 95CI | Upper 95CI | RR |
| --- | --- | --- | --- | --- | --- | --- | --- | --- |
| Time to knockdown | AIRID | Kisumu | 0.1 | 37.68 | 1.18 | 35.37 | 40.00 |  |
|  |  |  | 2 | 33.97 | 0.83 | 32.35 | 35.60 |  |
|  |  | Cove | 0.1 | Excluded from analysis. 50% effect threshold not reached |  |  |  |  |
|  |  |  | 2 | Excluded from analysis. 50% effect threshold not reached |  |  |  |  |
|  | KEMRI | Kisumu | 0.1 | 32.41 | 2.37 | 27.77 | 37.06 |  |
|  |  |  | 2 | 1.22 | 0.57 | 0.10 | 2.34 |  |
|  |  | Siaya | 0.1 | Excluded from analysis. 50% effect threshold not reached |  |  |  |  |
|  |  |  | 2 | 7.87 | 0.31 | 7.25 | 8.48 | 6.44 |
|  | LSTM | Kisumu | 0.1 | 13.24 | 1.08 | 11.12 | 15.36 |  |
|  |  |  | 2 | 5.64 | 0.21 | 5.23 | 6.06 |  |
|  |  | Siaya | 0.1 | Excluded from analysis. 50% effect threshold not reached |  |  |  |  |
|  |  |  | 2 | 4.08 | 0.78 | 2.55 | 5.62 | 0.63 |
|  |  | *Tiassale* | 0.1 | Excluded from analysis. 50% effect threshold not reached |  |  |  |  |
|  |  |  | 2 | 2.57 | 0.64 | 1.32 | 3.82 | 1.93 |
|  |  | *KDR* | 0.1 | Excluded from analysis. 50% effect threshold not reached |  |  |  |  |
|  |  |  | 2 | 7.86 | 0.48 | 6.92 | 8.80 | 1.38 |

For 60-minute knockdown at LSTM, the Kisumu reference strain had an EC□□KD of 0.052 mg/paper, 95% CI 0.034 to 0.071. Siaya showed a similar response, with an EC□□KD of 0.061 mg/paper, 95% CI −0.089 to 0.210, and an RR□□KD of 1.17. Tiassalé 13 showed a rightward shift, with an EC□□KD of 0.126 mg/paper, 95% CI −0.049 to 0.301, and an RR□□KD of 2.42. KDR showed the greatest rightward shift in knockdown response at LSTM, with an EC□□KD of 0.148 mg/paper, 95% CI −0.074 to 0.370, and an RR□□KD of 2.85. The confidence intervals for Siaya, Tiassalé 13, and KDR were wide and crossed zero, indicating low precision in these estimates, but the point estimates were consistent with reduced knockdown susceptibility in Tiassalé 13 and KDR relative to Kisumu.

At AIRID, the Kisumu reference strain had an EC□□KD of 0.007 mg/paper, 95% CI 0.004 to 0.009. Cové did not reach 50% knockdown within the tested dose range, so no EC□□KD estimate was reported for this strain.

At KEMRI, the Kisumu reference strain had an EC□□KD of 0.044 mg/paper, 95% CI 0.031 to 0.057. Siaya showed a clear rightward shift in knockdown response, with an EC□□KD of 0.148 mg/paper, 95% CI 0.111 to 0.185, and an RR□□KD of 3.36.

For 24-hour mortality at LSTM, the Kisumu reference strain had an EC□□Mort of 0.054 mg/paper, 95% CI 0.036 to 0.073. Siaya showed a comparable mortality response, with an EC□□Mort of 0.048 mg/paper, 95% CI 0.032 to 0.065, and an RR□□Mort of 0.89. Tiassalé 13 demonstrated markedly reduced susceptibility, with an EC□□Mort of 0.300 mg/paper, 95% CI 0.194 to 0.407, and an RR□□Mort of 5.56. KDR showed the greatest reduction in mortality response, with an EC□□Mort of 0.359 mg/paper, 95% CI 0.204 to 0.513, and an RR□□Mort of 6.65.

At AIRID, the Kisumu reference strain had an EC□□Mort of 0.030 mg/paper, 95% CI 0.020 to 0.039. Cové did not reach 50% mortality within the tested dose range, so no EC□□Mort estimate was reported. This was consistent with a high degree of reduced susceptibility under the conditions tested.

At KEMRI, the Kisumu reference strain had an EC□□Mort of 0.023 mg/paper, 95% CI 0.016 to 0.030. Siaya demonstrated reduced mortality susceptibility, with an EC□□Mort of 0.130 mg/paper, 95% CI 0.094 to 0.167, and an RR□□Mort of 5.65.

Time to 50% knockdown, TKD□□, was further estimated at each dose to characterise the rate of knockdown response across strains and sites. At LSTM, the Kisumu reference strain had a TKD□□ of 13.24 minutes, 95% CI 11.12 to 15.36, at 0.1 mg/paper, decreasing to 5.64 minutes, 95% CI 5.23 to 6.06, at 2 mg/paper. At 0.1 mg/paper, Siaya, Tiassalé 13, and KDR did not reach 50% knockdown within the observation period, so no TKD□□ estimates were reported at this dose. At 2 mg/paper, Siaya had a TKD□□ of 4.08 minutes, 95% CI 2.55 to 5.62, with an RR of 0.63. Tiassalé 13 had a TKD□□ of 2.57 minutes, 95% CI 1.32 to 3.82, with an RR of 1.93. KDR had a TKD□□ of 7.86 minutes, 95% CI 6.92 to 8.80, with an RR of

1.38. These results indicate that KDR showed slower knockdown than Kisumu at the highest dose, while the interpretation of Siaya and Tiassalé 13 should be treated cautiously because their lower-dose responses did not reach the 50% threshold (Table 2).

At AIRID, the Kisumu reference strain had a TKD□□ of 37.68 minutes, 95% CI 35.37 to 40.00, at 0.1 mg/paper, decreasing slightly to 33.97 minutes, 95% CI 32.35 to 35.60, at 2 mg/paper. This reflected the slower overall rate of knockdown at this site. Cové did not reach 50% knockdown at either dose, so no TKD□□ estimates were reported for this strain.

At KEMRI, the Kisumu reference strain had a TKD□□ of 32.41 minutes, 95% CI 27.77 to 37.06, at 0.1 mg/paper, decreasing to 1.22 minutes, 95% CI 0.10 to 2.34, at 2 mg/paper. Siaya did not reach 50% knockdown at 0.1 mg/paper, but at 2 mg/paper had a TKD□□ of 7.87 minutes, 95% CI 7.25 to 8.48, with an RR of 6.44, indicating slower knockdown relative to the Kisumu reference at the same dose.

### Applying the Two Tube Volatile Assay for routine susceptibility monitoring

Full validation of the Two Tube Volatile Assay has not yet been completed. The work presented here should therefore be considered an initial validation and use case dataset, demonstrating the method’s potential for measuring mosquito responses to controlled vapour exposure and providing a framework for further validation and implementation. Validation is being undertaken using the staged bioassay method validation framework described by Matope et al. (18), which defines preliminary development, feasibility testing, internal validation, and external validation as sequential stages in establishing whether a method is fit for purpose.

The preliminary development stage has been completed. The assay was designed to use standard WHO tube test hardware to generate controlled, non-contact exposure to transfluthrin vapour, while preventing mosquito contact with treated papers. The intended method scope was defined as a laboratory assay for measuring susceptibility to volatile insecticides, using knockdown during a 60-minute exposure and 24-hour mortality as the principal entomological endpoints. Negative controls, transfluthrin dose ranges, mosquito numbers, exposure duration, knockdown scoring intervals, and post-exposure holding conditions were defined, and these were incorporated into a draft SOP. Initial testing confirmed that the assay can generate concentration-dependent and time-dependent responses to transfluthrin and can distinguish between mosquito strains with different susceptibility phenotypes.

At LSTM, the assay was used to test multiple *Anopheles gambiae* strains, including the susceptible Kisumu reference strain and resistant or field-derived strains, across independent assay days. These experiments demonstrated repeatable concentration response relationships for both 60-minute knockdown and 24-hour mortality and showed that resistance ratios could be calculated relative to a susceptible reference strain tested under the same conditions.

Preliminary robustness testing has identified some practical considerations for routine use, particularly around airflow ((19)). Additionally, the need to control temperature, humidity, closely is also important as other studies have shown these influence transfluthrin volatilisation and vapour distribution within the test system, (20). However, formal acceptability criteria for within-day precision, between-day precision, operator effects, and environmental tolerance limits remain to be fully defined through continued testing.

This study represents an initial external validation through independent implementation of the assay developed at LSTM at AIRID and KEMRI. These sites used the same general assay format to test susceptible reference strains alongside resistant or field-derived mosquito strains. The results showed dose-dependent and time-dependent responses at each site and demonstrated clear separation between susceptible and less susceptible strains when analysed relative to the local susceptible reference. This provides preliminary evidence that the method can be transferred between laboratories and can detect reduced susceptibility in geographically and genetically distinct mosquito populations. However, these data do not yet constitute full external validation. Further work is required using a predefined external validation protocol, harmonised SOP, predefined performance criteria and a larger number of independent testing days and sites.

The remaining validation work should therefore focus on defining the analytical performance of the assay under routine use. This should include formal estimation of within-day repeatability, between-day precision, between-operator variation, and between-site reproducibility. A wider concentration range should be tested against highly resistant strains where 50% knockdown or mortality is not reached within the current dose range, so that the reportable range of the assay can be better defined. Additional testing should also assess the robustness of the method to controlled variation in key assay conditions, including temperature, humidity, airflow, mosquito age, mosquito density per tube, holding conditions and time between paper treatment and testing. These experiments will allow refinement of the SOP, confirmation of suitable dose ranges and development of a final method claim.

Based on the data generated in eight lab strains and three testing sites, we propose the following structured approach for adoption by the research and vector control community for routine monitoring of transfluthrin susceptibility in field populations. This framework defines how the assay can be applied, how data should be analysed and interpreted, and how additional characterisation can be undertaken where reduced susceptibility is detected.

Wider uptake and peer evaluation can be used to confirm reproducibility across laboratories, refine diagnostic concentrations, and establish harmonised guidance for transfluthrin susceptibility monitoring. As further data is generated the Two Tube Volatile Assay SOP can be expanded to additional species and volatile insecticides and applied using the same testing framework.

Routine susceptibility testing of a field population (**Error! Reference source not found.**):

- Test one negative control and three transfluthrin concentrations, 0.005, 0.1 and 2 mg/paper.
- Test the field population in parallel with a well characterised susceptible reference strain, of the same species where possible, under identical environmental conditions.
- Use three independent biological replicates, ideally across three separate assay days.
- Score knockdown during the 60-minute exposure at defined timepoints, for example 0, 5, 10, 15, 30, 45 and 60 minutes.
- Record 24-hour mortality as the primary endpoint.
- Fit dose response models to estimate EC□□KD and EC□□Mort for both populations.
- Calculate RR□□KD and RR□□Mort for the field population relative to the susceptible reference strain.
- Repeat testing at defined intervals, for example annually, to monitor changes over time.

**Interpretation of results**

- RRLJLJMort, based on 24-hour mortality, should be treated as the primary characterisation endpoint.
- RRLJLJKD provides a useful secondary endpoint and may reveal early or sublethal shifts not fully reflected in mortality.
- Comparison of RRLJLJMort and RRLJLJKD may help identify differences in response profile and may provide insight into underlying resistance mechanisms.
- Reductions in knockdown without corresponding shifts in mortality should be interpreted cautiously and confirmed with additional testing.
- All testing should be conducted under tightly standardised temperature, humidity, and airflow conditions to minimise variability in vapour availability and improve comparability across laboratories and timepoints.
- An increase in resistance ratios over time can be interpreted as evidence of reduced susceptibility.

If routine screening suggests reduced susceptibility, further characterisation should be undertaken by:

- generating a fuller concentration response curve using additional intermediate or higher doses,
- refitting the dose response model and re estimating ECLJLJ values, and
- recalculating resistance ratios relative to the susceptible reference strain tested under the same conditions.

If 50% knockdown or mortality is not reached within the standard screening range, this should not be treated as assay failure. Instead, it should be reported as failure to reach the 50% response threshold within the tested range, indicating that a wider or higher concentration range is required. This additional testing would allow more accurate quantification of the magnitude of reduced susceptibility and support longitudinal monitoring.

This proposed screening protocol is intended to be evaluated further in countries where spatial emanators are being assessed in trials or pilot implementation studies. Incorporating the Two Tube Volatile Assay alongside these studies would generate entomological susceptibility data from the same settings in which spatial emanator products are being evaluated. This would support interpretation of trial outcomes, provide baseline and follow-up susceptibility data for local vector populations, and help determine whether reduced susceptibility to transfluthrin is emerging over time. Data generated through these studies can also be used to refine diagnostic or monitoring concentrations, assess whether the same framework can be applied to other volatile active ingredients, and inform future guidance for susceptibility monitoring of transfluthrin and other spatial emanator products.

## Discussion

We have developed the Two Tube Volatile Assay and a protocol for routine susceptibility monitoring for the volatile pyrethroid transfluthrin. The assay is a modification of the WHO tube bioassay that enables controlled, non-contact vapour exposure to transfluthrin using standard WHO hardware, addressing a limitation in current susceptibility monitoring for spatial repellents. The WHO bottle bioassay exposes mosquitoes via both tarsal contact and exposure to vapour phases and is therefore unsuitable for isolating airborne effects. For volatile actives such as transfluthrin, where protection depends on spatial dispersion, a vapour-only assay is required to measure phenotypic response intrinsic to the active ingredient. Importantly, the assay measures toxicological effects of transfluthrin vapour exposure (knockdown and mortality), rather than behavioural repellence or avoidance responses.

Across all strains tested, transfluthrin produced clear concentration- and time-dependent knockdown and 24-hour mortality under vapour-only exposure. Most strains carrying resistance mechanisms against solid state pyrethrouids and other insecticides showed right-shifted response curves and elevated EC_50_ values relative to susceptible reference strains. Overall, the results align with known resistance profiles, where, for example, Tiassalé 13 carries multiple resistance mechanisms (15) and KDR carries primarily target site mutation (16). However, the magnitude of resistance observed to transfluthrin vapour was not always equivalent to resistance reported for contact pyrethroids. For example, KDR has previously been reported to show #14.6-fold resistance to deltamethrin relative to Kisumu (7), whereas the resistance ratios observed here for transfluthrin were lower. This suggests that resistance to contact pyrethroids does not necessarily translate directly into equivalent resistance to vapour-phase transfluthrin exposure.

This interpretation is supported by Zoh et al. (2023), who demonstrated through controlled selection experiments that deltamethrin and transfluthrin selected for distinct transcriptomic profiles in *Anopheles gambiae*, with minimal overlap in candidate resistance genes (21). Although the polyfluorinated structure of transfluthrin was initially thought to confer resilience to P450-mediated metabolism, selection with transfluthrin nevertheless upregulated specific P450 enzymes, suggesting that metabolic resistance to transfluthrin may be mechanistically distinct from resistance acting against contact pyrethroids. Together with the deli-pot data from Kokkas et al. (2026), these findings provide converging evidence that bioassay systems targeting the contact route alone may not reliably capture the full resistance phenotype relevant to volatile insecticide formulations (7).

The Siaya results also demonstrate why vapour-phase susceptibility should not be inferred directly from contact insecticide bioassays. Siaya was scored as resistant using WHO bottle contact bioassays, but in the LSTM Two Tube Volatile Assay its response was close to the Kisumu susceptible reference, particularly for 24-hour mortality. In contrast, the Siaya population tested at KEMRI showed clear rightward shifts relative to the local Kisumu reference. This difference between the two Siaya datasets may reflect colony history, local rearing conditions, differences in resistance intensity, or site-specific assay conditions. Further paired testing of Siaya colonies using harmonised replicate structures would be needed to determine whether these differences reflect biological variation between colonies or residual inter-site variation in assay performance.

The assay was conducted independently at three laboratories, LSTM, AIRID and KEMRI, enabling an initial assessment of inter-site reproducibility. Although absolute EC□□ estimates for Kisumu differed between sites, both knockdown and mortality values were consistent with Kisumu behaving as a susceptible reference strain at each laboratory. For 60-minute knockdown, Kisumu EC□□KD values were 0.052 mg/paper at LSTM, 0.007 mg/paper at AIRID and 0.044 mg/paper at KEMRI. For 24-hour mortality, Kisumu EC□□Mort values were 0.054 mg/paper at LSTM, 0.030 mg/paper at AIRID and 0.023 mg/paper at KEMRI. These differences in absolute estimates are likely to reflect normal between-site variation in assay conditions, including temperature, airflow speed, and directionality, as well as some subjectivity in knockdown scoring, but it’s also possible there is a meaningful difference in biological susceptibility of the reference strain.

Testing with transfluthrin is likely to be particularly sensitive to environmental and procedural variation because exposure depends on volatilisation and movement of the active ingredient through the test system. The generation of resistance ratios relative to a susceptible reference strain tested in parallel is therefore proposed to make the method less sensitive to variability between sites or testing days. This was evident at KEMRI, where Kisumu EC□□ values of 0.044 mg/paper for knockdown and 0.023 mg/paper for mortality were obtained, while Siaya showed clear rightward shifts in response, with an RR□□KD of 3.36 and an RR□□Mort of 5.65. This indicates that the assay can be implemented across laboratories using standard WHO test kits and that susceptibility classification is robust to the level of variation observed between sites.

Testing at AIRID additionally included the resistant Covè strain from Benin. Covè did not reach 50% knockdown or 50% mortality within the tested dose range, so EC□□KD, EC□□Mort and RR□□ estimates were not reported from the current analysis. This result is still informative, as failure to reach the 50% response threshold even at 2 mg/paper indicates reduced susceptibility relative to the AIRID Kisumu reference. However, higher concentrations would be required to fit a complete concentration-response curve and fully characterise the magnitude of resistance in this strain. It is recommended that when performing susceptibility monitoring against a population that higher concentrations are tested where the response with the standard concentrations does not reach ∼100%.

These findings for the Covè strain are at odds with those reported by N’dombidjé et al. (2026), who evaluated Mosquito Shield™ in experimental huts against wild, free-flying, pyrethroid-resistant *Anopheles gambiae s.l*. at the Covè field station in southern Benin over two 32-day product life cycles. Despite elevated levels of pyrethroid resistance in the local vector population, Mosquito Shield™ achieved 43% protective efficacy against landing, 64% personal protection against blood-feeding, and induced 49% mortality. Together, these findings suggest that although substantial reductions in susceptibility to transfluthrin vapour can be detected under controlled laboratory conditions, transfluthrin-based spatial emanators may still retain meaningful entomological efficacy in operational settings where conventional pyrethroids perform poorly (22).

The AIRID Covè results are also notable when considered alongside the findings of Okeyo et al. (2025), where the highly resistant Pimperena strain required more than 1000-fold higher transfluthrin doses than the Kisumu susceptible strain to produce comparable responses.

While the resistance observed in Covè was not quantified to this extent within the tested dose range, the inability to reach 50% knockdown or mortality at the highest concentration tested suggests that similarly high-intensity resistance phenotypes may occur in some populations exposed to volatile pyrethroids (23).

KEMRI additionally tested the Siaya field strain, which showed a knockdown resistance ratio (RR□□KD) of 3.36 and a mortality resistance ratio (RR□□Mort) of 5.65 relative to the local Kisumu reference strain. This was consistent with reduced susceptibility in Siaya under the KEMRI assay conditions, particularly for the mortality endpoint. The difference between the two Siaya datasets may partly reflect divergence in resistance allele frequencies between the two colonies over time. Okeyo et al. (2025) found that reduced behavioural sensitivity to transfluthrin in *An. gambiae* from western Kenya was associated with overexpression of detoxification genes, particularly CYP12F2, alongside high frequencies of kdr L995S the same allele documented in Siaya. Genotypic characterisation of both Siaya colonies at the time of testing would help to determine whether observed differences in phenotypic response correspond to differences in underlying resistance mechanism frequencies (23). Together, the multi-site data indicate that the method is capable of detecting reduced susceptibility in geographically and genetically distinct field-derived strains beyond those used in initial development, and that results are broadly reproducible across independent laboratories when interpreted relative to a susceptible reference strain tested in parallel. A discriminating concentration assay is designed to detect the early signs of emerging resistance to an insecticide, using mortality against a single dose in a bottle or tube to classify a mosquito population as susceptible or resistant (5,17). Such an approach is suited to surveillance in previously susceptible populations where the primary goal is an early warning flag for the emergence of resistance. In contrast, the Two Tube Volatile Assay is designed to characterise the response of mosquitoes to transfluthrin in populations where pyrethroid resistance mechanisms are likely to already be present. Rather than applying a binary susceptible/resistant classification, this assay generates a full concentration-response curve from which a resistance ratio can be calculated relative to a susceptible laboratory reference colony. Monitoring for an increase in this ratio over time would indicate that transfluthrin-specific resistance is developing or intensifying within the target population, providing a quantitative tool for longitudinal surveillance in settings with existing resistance (7).

Overall, the Two Tube Volatile Assay reliably distinguished susceptible and resistant laboratory strains and quantified distinct levels of susceptibility between strains with different resistance mechanisms. Resistance ratios were more pronounced for mortality than knockdown in the strongly resistant strains, suggesting that mortality may provide a more stable endpoint for classification, though confidence intervals for some estimates (particularly Covè mortality and Tiassalé knockdown) were wide, reflecting the difficulty of fitting complete concentration-response curves for all strains within the tested range.

Imprecision in some knockdown estimates highlights the importance of adequate dose spacing and replication when generating full concentration response curves. By measuring resistance ratios compared to a susceptible reference strain using both endpoints, and by repeat testing over time, we propose the assay and monitoring protocol developed here to be suitable to characterise the response to transfluthrin in a population being targeted with transfluthrin-based spatial emanators, to detect changes in susceptibility over time, and to provide some evidence about possible resistance mechanisms which can be further explored with, for example, molecular analysis, as required.

Limitations of the assay should be recognised, and further development and validation is required. First, the method is validated here for transfluthrin due to the urgent need for new methods for monitoring susceptibility to this active ingredient. Application to other volatile actives will require compound specific optimisation and validation. Similarly, we present data for *Anopheles gambiae* as a key malaria vector targeted by spatial emanators, but further testing is required to determine efficacy of the method and proposed transfluthrin concentrations to other vector species. Second, mass of transfluthrin applied to a filter paper only serves as a proxy for exposure, as vapour concentration in the tube is not directly quantified. Environmental factors such as temperature, humidity and airflow influence volatilisation and must be tightly standardised to ensure comparability between assays. It is important to note when interpreting the results of transfluthrin susceptibility testing that the purpose of the Two Tube Volatile Assay is controlled comparative susceptibility testing of the target mosquitoes rather than replication of operational exposure scenarios to evaluate a product or predict field efficacy. The confined tube space does not aim to simulate field dispersion. We are not aiming to predict product efficacy with this assay, but to characterise susceptibility and monitor for changes in response. Reduced response to transfluthrin observed in this laboratory format does not necessarily imply loss of operational effectiveness of spatial emanators in field settings. Reduced intrinsic susceptibility measured under confined laboratory vapour exposure reflects one component of the interaction between mosquito and active ingredient, but operational impact depends on transmission dynamics within shared spaces.

Even where mortality or knockdown responses are reduced in target populations, spatial emanators may continue to reduce disease transmission through altered host seeking behaviour, biting distribution, and resulting exposure risk at the community level, as indicated by experimental hut results in Cové. Spatial emanators are not purely personal protection tools. Unlike treated nets, which primarily protect the individual user, airborne actives may confer community level effects by diverting or suppressing host seeking within shared airspace. Even if some individuals exhibit reduced susceptibility, diverted mosquitoes may rebound to alternative hosts, altering but not necessarily eliminating population level protection dynamics. This redistribution effect is distinct from purely individual protection. A reduction in susceptibility may change where bites occur but does not automatically eliminate suppression at the population scale. Epidemiological impact depends on transmission networks rather than individual mortality alone (24). By collecting susceptibility data using the Two Tube Volatile Assay alongside measuring SE efficacy in different sites we can hope to explore the relationship between the two. This new assay will allow a change in transfluthrin toxicity against a target field population, but additional methods could potentially be used to characterise and detect any changes in other endpoints such as blood feeding inhibition or repellence.

The distinction between spatial emanators and insecticide-treated nets is also relevant to how reduced susceptibility should be interpreted and monitored. Treated nets create an exposure scenario tied to a specific behaviour, requiring consistent nightly use and creating gaps in protection when individuals are awake, when they do not sleep under the net, or when they rise during the night, periods during which exposure risk may be substantial (25,26). In contrast, spatial emanators operate through continuous passive airborne emission over shared airspace, with dose exposure determined by proximity and air circulation rather than user behaviour. This difference in exposure route and variability has methodological implications for susceptibility testing. Reduced epidemiological effects in net-based products can be confounded by variation in net use behaviour, which itself responds to perceived disease risk, making it difficult to separate intrinsic susceptibility of mosquitoes from usage dynamics, and to interpret entomological susceptibility data alongside epidemiological outcomes. Spatial emanators, by contrast, do not depend on behavioural decisions at the point of exposure, meaning susceptibility monitoring reflects the intrinsic response to the active ingredient more directly and may therefore be more relevant to field exposure.

However, mortality represents only one measurable endpoint within a broader behavioural cascade. Spatial repellents potentially act through knockdown, excito-repellence, flight disruption, feeding inhibition, and diversion (20). Blood-feeding inhibition as an additional mode of action is now well established, with the disruption of host-seeking and feeding behaviour leading to reduced mosquito landing and blood-feeding success (24,27,28). This blood-feeding inhibition, alongside disarming effects, underpins their ability to reduce human–vector contact and transmission risk in both semi-field and field settings. Laboratory mortality resistance ratios quantify only part of the phenotype. Even where mortality susceptibility is reduced, sublethal behavioural disruption may still suppress biting or delay feeding sufficiently to reduce transmission potential. Resistance classification based solely on knockdown and mortality risks oversimplifying predictions about operational impact and data should be interpreted and used within this context.

A limitation of the present multi-site comparison is that replicate structure was not fully balanced across sites. While LSTM replicates were distributed across three assay days, KEMRI and AIRID replicates were conducted within a single day. As a result, the KEMRI and AIRID data provide evidence of within-day repeatability and successful implementation of the method at independent testing sites, but they do not independently quantify between day variability. Future applications of the assay should therefore use the recommended three-day replicate structure to support more robust comparisons between strains, sites, or time points.

A range of methods already exists for measuring the entomological or behavioural effects of spatial emanators, but these are oriented toward efficacy rather than susceptibility testing of the active ingredient. Large testing arenas such as the Peet-Grady chamber or a two-room test can quantify entomological effect (13) but are impractical for routine or field-based susceptibility testing and do not lend themselves to characterising the response of mosquitoes to the active ingredient itself. Other WHO-recommended methods for assessing responses to volatile compounds, including the Y-tube olfactometer and the HIITSS chamber, are benchtop-scale but require custom-built equipment and tightly controlled experimental conditions (13,14). While these systems can be standardised, they are not as routinely implemented or widely accessible as established WHO tube tests or CDC bottle assays, limiting their suitability for routine or large-scale resistance monitoring, particularly at field stations where most susceptibility testing is conducted. These methods address efficacy testing, but a simple, scalable, non-contact method for susceptibility monitoring remained a gap, which the Two Tube Volatile Assay is designed to fill.

Despite these constraints and the further validation required, the Two Tube Volatile Assay already offers clear advantages over existing testing methods for transfluthrin susceptibility monitoring. Unlike the WHO bottle bioassay, which combines tarsal contact and vapour exposure and provides a binary classification at a single discriminating concentration, the Two Tube Volatile Assay isolates vapour-phase exposure and generates concentration response data. This allowed strains classified as resistant in the bottle bioassay to be separated by their response profiles. For example, Siaya, KDR and Tiassalé 13 were all classified as resistant in the bottle bioassay, but the Two Tube Volatile Assay showed that LSTM Siaya responded similarly to Kisumu, while KDR and Tiassalé 13 showed clear reductions in mortality and elevated resistance ratios. Cové showed an even stronger reduction in susceptibility, with limited knockdown and mortality even at 2 mg/paper. The assay therefore provides information on dose dependency, knockdown dynamics, 24-hour mortality and resistance ratios relative to a susceptible comparator, while avoiding the confounding effect of direct contact exposure. Because operational exposure to transfluthrin from spatial emanators is primarily airborne, this vapour-only format is better suited for characterising relevant susceptibility phenotypes and monitoring changes over time. The method is simple, inexpensive, based on widely available WHO tube test kits, and provides a scalable platform for evaluating susceptibility to transfluthrin and other volatile active ingredients within its intended scope.

Future work should focus on measuring inter-laboratory reproducibility, further refinement of recommended concentrations if indicated by testing with further mosquito populations, integration with quantitative air sampling to strengthen interpretation of the response to exposure, and determination of whether a single discriminating concentration derived from the concentration-response data presented here is warranted and sufficient for routine surveillance purposes. Further validation with additional field and laboratory populations will be required to assess this. Complementary behavioural assays in larger arenas would also help to characterise sublethal effects on host seeking and feeding. By quantifying the within-and between-day variability in results from the assay we will be able to conduct power calculations to refine the protocol and interpretation of the results, as has been done for other standard bioassays (29–32).

## Conclusion

The Two Tube Volatile Assay provides a practical benchtop method for monitoring susceptibility to transfluthrin vapour. By preventing contact exposure while retaining standard WHO test kit, it enables controlled comparison of intrinsic responses to a volatile pyrethroid using a common and scalable platform.

Across 5 colonies of *An. gambiae* with resistance to contact insecticides tested in three independent labs, and susceptible reference strains maintained in each lab, the assay differentiated susceptible and resistant laboratory strains and generated clear concentration dependent responses. Based on these data, 24-hour mortality is recommended as the primary endpoint for routine resistance monitoring using this method. Sixty-minute knockdown provides a useful secondary endpoint, particularly for detecting early or sublethal shifts in response that may not yet translate into major differences in mortality. The metrics for characterising the response to transfluthrin are mortality resistance ratio (RR□□Mort) and knockdown resistance ratio (RR_50KD_), compared to a susceptible reference strain.

For routine susceptibility monitoring, we propose a three concentration plus control testing scheme. Results should be compared directly against data generated for a known susceptible reference strain of the same species under identical conditions. Where reduced mortality is observed, extended characterisation using a fuller concentration response curve can be undertaken to estimate EC_50_ more accurately and calculate RR_50_ values. Dedicated behavioural assays would be required to characterise sublethal effects such as blood feeding inhibition independently of knockdown and mortality and would provide more information about the response of a population to transfluthrin and specific mechanisms of resistance.

Based on the observed performance of the assay and to improve robustness for future susceptibility monitoring, we recommend that assays are conducted as three independent biological replicates performed across three separate days, with one replicate per dose per day where feasible. This design captures both within-assay and between day variation and reduces the risk that site-specific environmental or colony handling effects on a single day disproportionately influence estimated RR□□Mort and RR□□KD values.

The work reported here constitutes the preliminary development stage of a structured method validation process, following the framework proposed by Matope et al. (2023) for bioassay methods used to evaluate vector control tools (18). Within this stage, method design, endpoints, and key testing conditions were defined, and baseline experiments were conducted. Multiple robustness sub-studies are planned to examine variables including mosquito number per tube, sample orientation, and solution stability over time, to identify sources of variability and build robustness into the method prior to formal validation.

Provisional acceptability criteria for the primary endpoints of 24-hour mortality and 60-minute knockdown will be defined based on the results of these experiments, accounting for the inherent biological variability of the test system.

The next planned stage is feasibility experimentation, in which within-day and between-day precision will be formally estimated. These data will underpin a formal sample size calculation for internal validation, in which the analytical performance of the method will be tested within a single laboratory against pre-specified acceptability criteria, and a draft standard operating procedure (SOP) will be produced. Subsequent external validation across at least two independent laboratories will be required to fully establish reproducibility of the method claim and support its wider implementation for routine volatile pyrethroid susceptibility monitoring.

Within its intended scope, the assay offers a simple, standardisable approach for volatile pyrethroid susceptibility monitoring and provides a foundation for the development of harmonised testing guidance for transfluthrin and other volatile compounds.

## Declarations

### Ethical approval and consent

Ethical approval and consent were not required.

### Data availability

#### Underlying data

Zenodo: Two Tube Volatile Assay Dataset, https://doi.org/10.5281/zenodo.19370203 (Praulins et al., 2026).

This project contains the following underlying data:

- Two_Tube_Time_Course_Data.xlsx
- Bottle_Bioassay_Data.xlsx

Data are available under the terms of the Creative Commons Attribution 4.0 International license (CC-BY 4.0) (https://creativecommons.org/licenses/by/4.0/).

#### Software availability

Source code available from: https://github.com/GioP93/Two Tube -Volatile-Bioassay

Archived source code is available from https://doi.org/10.5281/zenodo.19370203 (Praulins et al., 2026).

Source code is available under the terms of the GNU General Public License version 3 (GPL-3.0-only) (https://opensource.org/license/gpl-3-0).

#### Author contributions

Giorgio Praulins – Conceptualization, Methodology, Data Curation, Formal Analysis, Visualization, Writing – Original Draft Preparation, Writing – Review & Editing, Project Administration.

Amy Lewis – Investigation, Methodology, Validation, Writing – Review & Editing. Tim Hill – Investigation, Validation, Writing – Review & Editing.

Boris N’dombidje - Investigation, Writing – Review & Editing.

Gemma Harvey – Investigation, Writing – Review & Editing, Data Curation, Formal Analysis Daniel McDermott – Conceptualization, Methodology, Writing – Review & Editing.

Corine Ngufor - Funding Acquisition, Supervision, Writing – Review & Editing.

Rosemary Susan Lees – Conceptualization, Funding Acquisition, Supervision, Writing – Review & Editing, Project Administration

